# Angiotensin converting enzyme 2 (ACE2): Virus accomplice or host defender

**DOI:** 10.1101/2022.03.06.483197

**Authors:** Jiyan Wang, Hongkai Chang, Yaya Qiao, Huanran Sun, Xichuan Li, Shuofeng Yuan, Shuai Zhang, Changliang Shan

## Abstract

The current coronavirus disease-19 (COVID-19) caused by the acute respiratory syndrome coronavirus 2 (SARS-CoV-2) infection has seriously disrupted the daily life of human, mainly attributed to the fact that we know too little about SARS-CoV-2. Increasing studies show that viral infection alters host cells glucose metabolism, which is crucial for viral nucleic acid replication. Here, we integrated RNA-sequencing results and found that SARS-CoV-2 infection alters the aerobic glycolysis, pentose phosphate pathway (oxiPPP), and DNA replication in lung tissues and cells. However, the direction of metabolic flux and DNA replication were dominated by angiotensin-converting enzyme 2 (ACE2), a host cell-expressed viral receptor protein. More interesting, although hosts with high expression of ACE2 are more likely to be infected with SARS-CoV-2, the invading virus cannot perform nucleic acid replication well due to the restriction of glucose metabolism, and eventually resulting prolonged infection-cycle or infection failure. Our findings, after a typical epidemiological investigation and modeling analysis, preliminarily explain the reasons for the emergence of asymptomatic infections or lower copy virus at early stage in host with higher ACE2 levels, which will provide important help for the development of more accurate and effective detection methods for diagnosing COVID-19.

**Graphical abstract:** 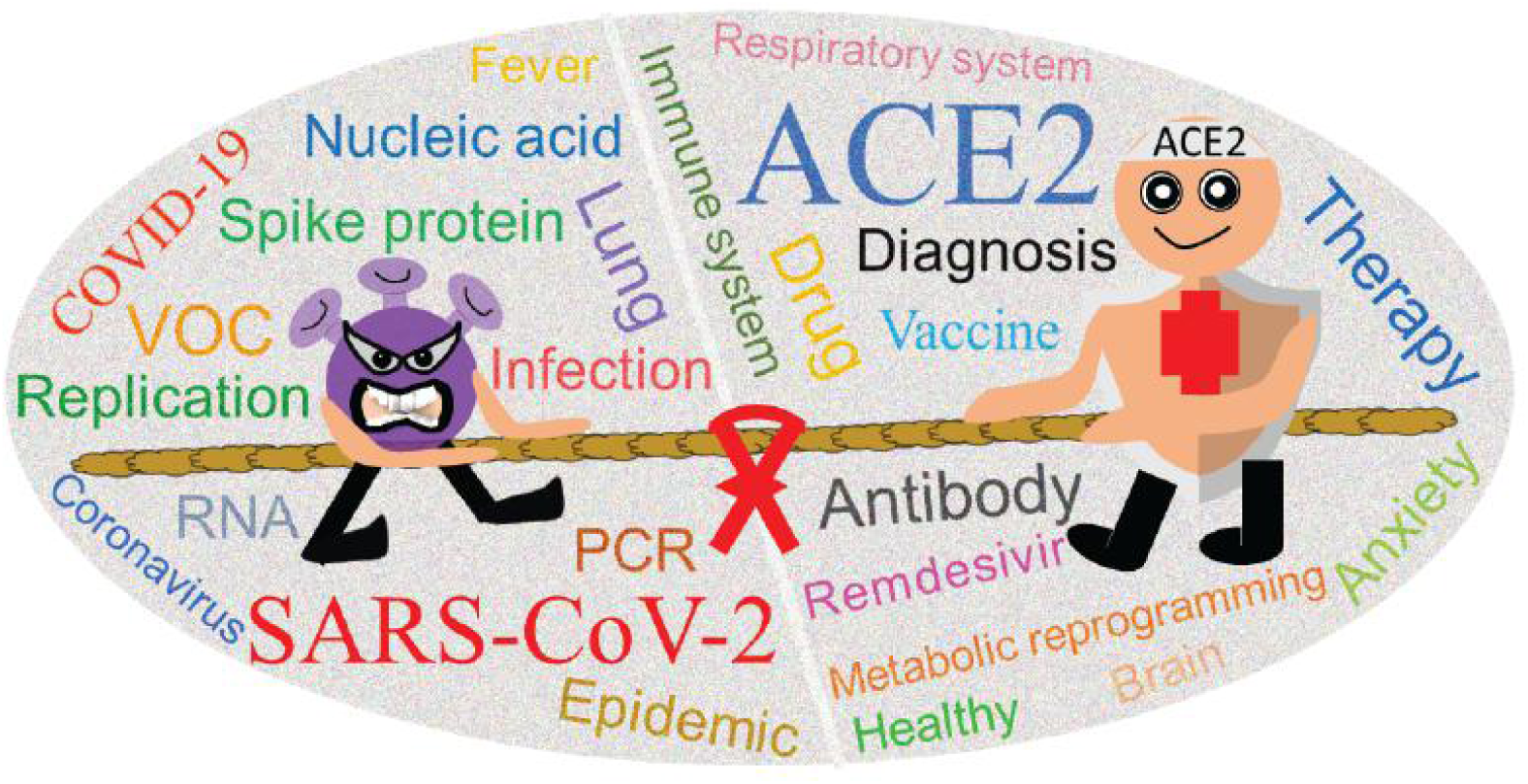

## INTRODUCTION

Since the end of 2019,^1^ the coronavirus disease-19 (COVID-19) caused by the acute respiratory syndrome coronavirus 2 (SARS-CoV-2) virus infection has swept the world, and has brought considerable effects to countries and people around the world. That’s how COVID-19 got its name. SARS-CoV-2 is a single, positive-stranded RNA virus enveloped in a lipid bilayer.^2,3^ The lipid bilayer fuses with the host cell membrane, releasing RNA into the cytoplasm and causing translation of various viral proteins. The replicated RNA genome and synthesized viral proteins reassemble into new viruses, which burst out of the cell.^4,5^ The virus enters host cells via interaction of two proteins. The viral counterpart is the spike-protein (S-protein), a glycoprotein expressed as a homotrimer on the viral envelope.^6^ This viral S-protein binds with the human protein receptor angiotensin converting enzyme 2 (ACE2).^7,8^ ACE2, as a trans-membrane protein, is abundant in lung, heart, kidney, and adipose tissue.^9,10^ Binding of S-protein with ACE2 allows for membrane fusion and entry of COVID-19 RNA into the cell. Therefore, the binding of two proteins serves as a target for potential treatments and vaccinations^11^. However, it has not been reported whether ACE2 has other roles in the process of SARS-CoV-2 infection and viral nucleic acid replication.

Compared to SARS, SARS-CoV-2 uses the same mechanism for entering host cells, but at slower speeds. However, SARS-CoV-2 accumulates more in the system, suggesting a stronger replication capacity of SARS-CoV-2. SARS-CoV-2 infection has an incubation period with an average of 3-9 days,^12–16^ which indicates that a person is contagious before symptoms appear, and this poses great difficulties for the prevention against COVID-19 epidemic. About 18% of cases are reported to be asymptomatic.^17–19^ Interestingly, younger patients tend to remain asymptomatic (even if constantly around an infected individual), while the elderly usually show symptoms,^12,18^ which implies that the virus infection is not absolutely successful. Besides, About 44% of transmissions have been reported to occur before symptoms appear,^20^ suggesting that SARS-CoV-2 is likely to replicate a large number of progeny viruses during the incubation period. Mild and asymptomatic cases tend to shed 10 days (between 8-15 days) after symptom resolution,^21,22^ with 90 % resolving after 10 days and nearly all cases resolving after 15 days.^21^ Therefore, restricting the movement of people and maintaining home isolation (or centralized isolation) are currently effective means of fighting against COVID-19 epidemic, which also hinder life and economic development simultaneously.

Reverse transcriptase-polymerase chain reaction (RT-PCR) remains the gold standard for diagnosing COVID-19.^23^ However, for asymptomatic infected persons, the presence of SARS-CoV-2 cannot be effectively detected, even by continuous detection. Although researchers around the world are currently studying COVID-19, we are still short of a comprehensive understanding for SARS-CoV-2. In this study, we found that ACE2, the host channel for virus entry, also inhibits viral nucleic acid replication. Specifically, the expression of highly expressed ACE2 is inhibited after infection with SARS-CoV-2, resulting in glucose metabolism, oxidative pentose phosphate pathway (oxiPPP), and DNA replication process inhibited, and ultimately inhibiting the replication of coronavirus. Although high ACE2 expression represents greater susceptibility, it does not mean that more progeny viruses will be produced. So, our analysis results suggested that the host with high ACE2 expression has a certain inhibitory effect on virus replication. Combined with epidemiological investigations and modeling analysis, we preliminarily explain the causes of asymptomatic infections, which will provide theoretical help for the development of more accurate methods for identifying positive infections.

## RESULTS

### Glucose metabolism is reprogramed in host infected of SARS-CoV-2

To ensure optimal environments for their replication and spread, viruses have evolved to alter many host cell pathways. Most viruses examined to date induce aerobic glycolysis also known as the Warburg effect.^24,25^ Consistent with previous reports,^26^ we analyzed the RNA-sequencing results in the published article,^27^ and found that the glucose metabolism was promoted in mice lung tissue infected SARS-CoV-2 (Fig. 1a). To further confirm these findings, we also analyzed in SARS-CoV-2 infected human normal bronchial epithelial cells (NHBE) and lung cancer cells (A549), and found that the glucose metabolism was also promoted by SARS-CoV-2 infection (Fig.1a).^28^ Unexpectedly, the glucose metabolism in ACE2-overexpressing A549 (A549-ACE2) cells and Calu-3 cells were blocked by SARS-CoV-2 infection (Fig. 1b). In addition to glucose metabolism, other metabolic processes, including tricarboxylic acid cycle (TCA cycle), are also regulated by SARS-CoV-2 infection, but glucose metabolism is the only pathway enriched in all SARS-CoV-2 infected cells. Although the direction of glucose metabolism was not identical in host cells with virus infection, our finding still suggested that glucose metabolism was significantly reprogrammed in host cells with SARS-CoV-2 infection.

**Fig. 1.**
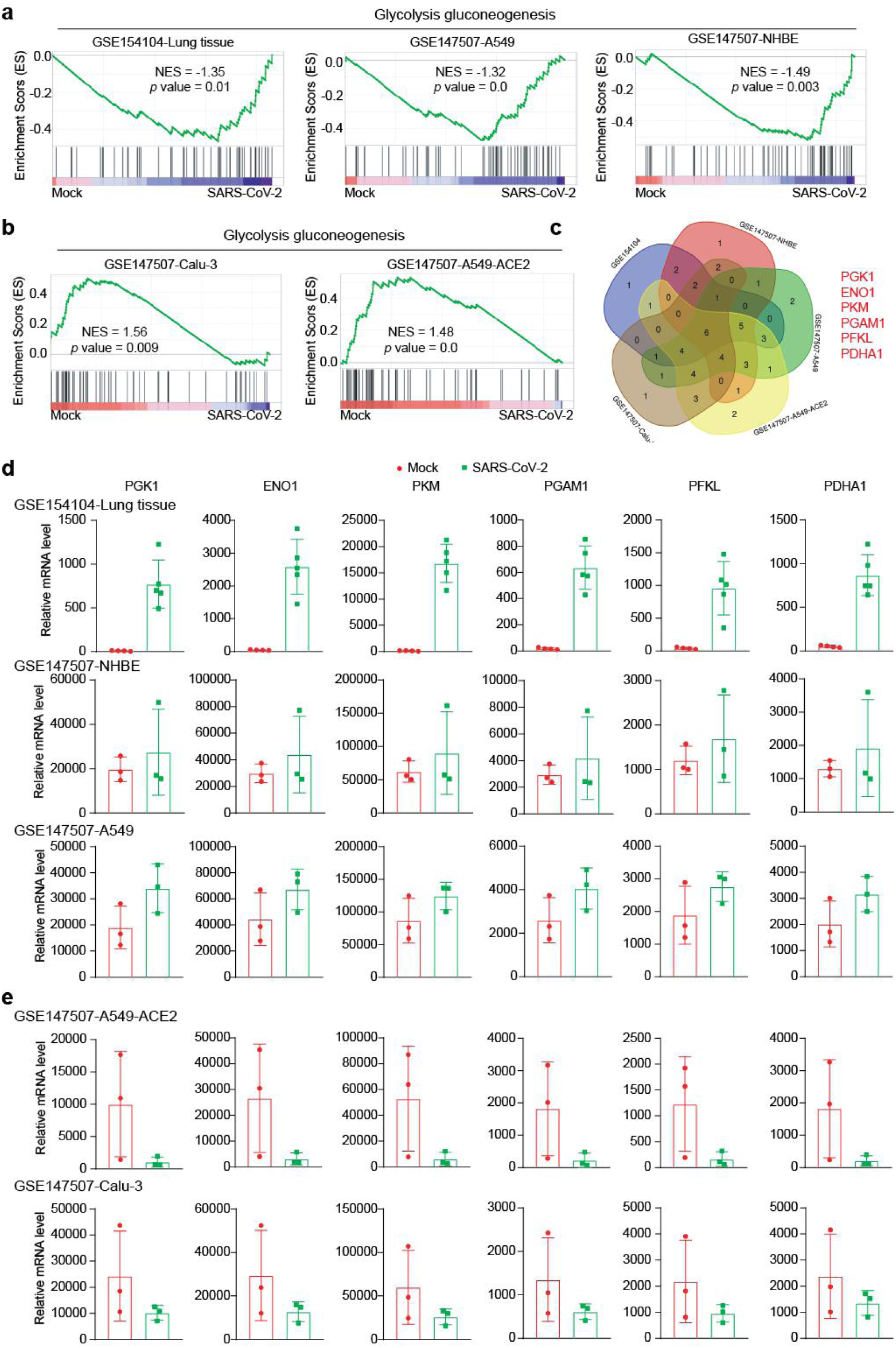
SARS-CoV-2 infection alters host glucose metabolism. **a,b** Gene set enrichment analysis (GSEA) pathway enrichment analyses of SARS-CoV-2 infection in mice lung tissue, lung normal cells, and lung cancer cells from the GEO datasets (GSE154104 and GSE147507). **c** The overlapping analysis of differentially expressed metabolic enzymes in mice lung tissue, lung normal cells, and lung cancer cells infected with SARS-CoV-2 from the GEO datasets (GSE154104 and GSE147507). **d, e** Expression changes of glucose metabolic enzymes in mice lung tissue, lung normal cells, and lung cancer cells infected with SARS-CoV-2 from the GEO datasets (GSE154104 and GSE147507).

Next, to screen out the key factors that are precisely regulated by virus, we performed an overlap analysis and found that six metabolic enzymes (PGK1, ENO1, PKM, PGAM1, PFKL and PDHA1) were identified as potential target (Fig. 1c). Indeed, when we analyzed the expression changes of these six metabolic enzymes after being infected by SARS-CoV-2 in different tissues and cells, and found that the expression of metabolic enzymes were up-regulated in mice lung tissue, NHBE cells, and A549 cells (Fig. 1d); while the expressions of metabolic enzymes were all down-regulated in A549-ACE2 and Calu-3 cells (Fig. 1e). These analysis results further confirm that SARS-CoV-2 infection reprograms the host’s glucose metabolism by increasing or decreasing the expression of metabolic enzymes. Here, we will explore the underlying mechanism about the promotion or inhibition of glucose metabolism by SARS-CoV-2 infection in the different host cells.

### DNA replication and oxidative pentose phosphate pathway are remodeled in host infected of SARS-CoV-2

Viral infection has been reported to remodel host’s glucose metabolism,^24^ and our above analysis also support this conclusion. In addition, glucose metabolism is closely related to viral infection and replication processes.^26,29^ More importantly, interfering with glucose metabolism inhibits the replication of SARS-CoV-2.^30^ Diabetes is a good model for the study of glucose metabolism, with an increased incidence and severity of SARS-CoV-2 infection in diabetic patients due to an imbalance in glucose metabolism,^31^ and diabetics with better controlled blood glucose have a higher survival rate.^32^ Meanwhile, infection with SARS-CoV-2 also increases the risk of diabetes,^33^ which indicates that the glucose metabolism pathway is hijacked by SARS-CoV-2. Furthermore, among the glucose metabolic enzymes that we identified by analyzing RNA-sequence data, PGK1 and ENO1 have been reported to have functions in promoting DNA replication and viral nucleic acid replication.^34,35^ The replication of SARS-CoV-2 also requires raw materials and energy from the host DNA replication process.

To this end, we analyzed DNA replication in tissues and cells infected with SARS-CoV-2. As shown in Fig. 2, the DNA replication was promoted in mice lung tissue and A549 cells with SARS-CoV-2 infection (Fig. 2a). While, the DNA replication was inhibited in A549-ACE2 and Calu-3 cells with SARS-CoV-2 infection (Fig. 2b). Thus, the DNA replication was tightly correlated with glucose metabolism reprogram in cells with SARS-CoV-2 infection. Based on the purpose of finding the target, we screened out 25 potential targets (Fig. 2c), which are important proteins during DNA replication. Next, we chose three proteins (RPA1, POLA1 and LIG1) that play a vital role in DNA replication among these targets, and found that their expression was also regulated by SARS-CoV-2 infection (Fig. 2d). These results indicate that SARS-CoV-2 not only reprograms glucose metabolism, but also acts on DNA replication process. Collectively, our analysis findings suggested that SARS-CoV-2 infection hijack host cells glucose metabolism to promote viral nucleic acid replication.

**Fig. 2.**
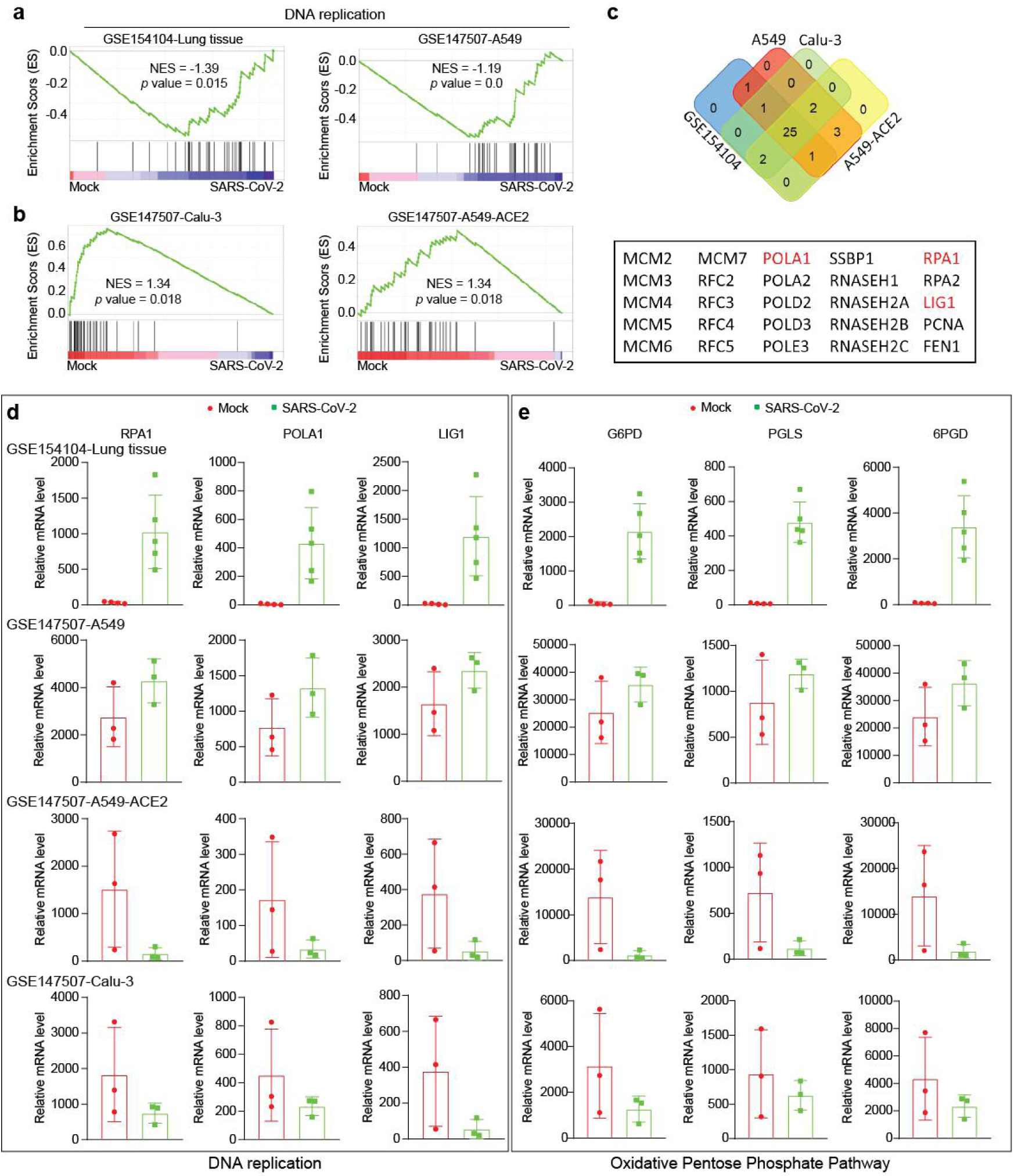
SARS-CoV-2 infection alters host DNA replication and oxidative pentose phosphate pathway (oxiPPP). **a, b** GSEA pathway enrichment analyses of SARS-CoV-2 infection in mice lung tissue and lung cancer cells from the GEO datasets (GSE154104 and GSE147507). **c** The overlapping analysis of differentially expressed DNA replication genes in mice lung tissue and lung cancer cells infected with SARS-CoV-2 from the GEO datasets (GSE154104 and GSE147507). **d, e** Expression changes of DNA replication and oxiPPP related genes in mice lung tissue and lung cancer cells infected with SARS-CoV-2 from the GEO datasets (GSE154104 and GSE147507).

As we know, glucose metabolism, especially for glycolysis mainly provides cells with energy and raw materials for biogenesis. Thus, the consistent changes in glucose metabolism and DNA replication suggest a possible connection between two pathways. The oxidative pentose phosphate pathway (oxiPPP) is a major glucose catabolic pathway that links glucose metabolism to the nucleotide synthesis. As a branch of glycolysis, the oxiPPP converts glucose-6-phosphate (G-6-P) to ribulose-5-phosphate (Ru-5-P), providing raw materials for nucleotide synthesis. Ribose, the basic component of nucleotides, provides the backbone for the synthesis of nucleotides. Due to ribose cannot be ingested through food, the oxiPPP is the main pathway for ribose synthesis. For coronaviruses, viral replication depends on the availability of cellular nucleotide pools,^36^ and compounds that inhibits nucleotide synthesis (such as ribavirin) has been found to inhibit the replication of SARS-CoV-2.^30^ In oxiPPP, G6PD, PGLS, and 6PGD are responsible for converting G-6-P to Ru-5-P.^37,38^ In the above-mentioned mouse lung tissue and lung cancer cells, we found oxiPPP enzymes (G6PD, PGLS and 6PGD) were regulated in the host with SARS-CoV-2 infection (Fig. 2e). The identical changes in glucose metabolism, oxiPPP, and DNA replication process show that metabolic reprogramming resulting from SARS-CoV-2 infection is cross-linked. After SARS-CoV-2 infection, the host’s metabolic system is hijacked to promote virus nucleic acid replication and reproduction.

### ACE2 determines how virus alters host’s metabolic balance

Although integrative analysis of RNA-sequencing results from published papers, and found that SARS-CoV-2 infection reprograms host glucose metabolism and oxiPPP to promote viral nucleic acid replication, however, it is surprising that changes in glucose metabolism and oxiPPP are difference in A549 and A549-ACE2 cell lines with different ACE2 levels (Fig. 1 and 2). Thus, we speculate that ACE2 may play an important role in determining these process. Meanwhile, it has been reported that the expression of ACE2 in Calu-3 cells is higher than that in A549 cells.^39^ To test our hypothesis, we compared the expression of ACE2 among NHBE, A549, A549-ACE2, and Calu-3 cells. The results showed that the expression in A549-ACE2 and Calu-3 cells was significantly higher than that in NHBE and A549 cells (Fig. 3a), which suggests that the expression level of ACE2 is a key point for determining the direction of host glucose metabolism, oxiPPP, and DNA replication.

**Fig. 3.**
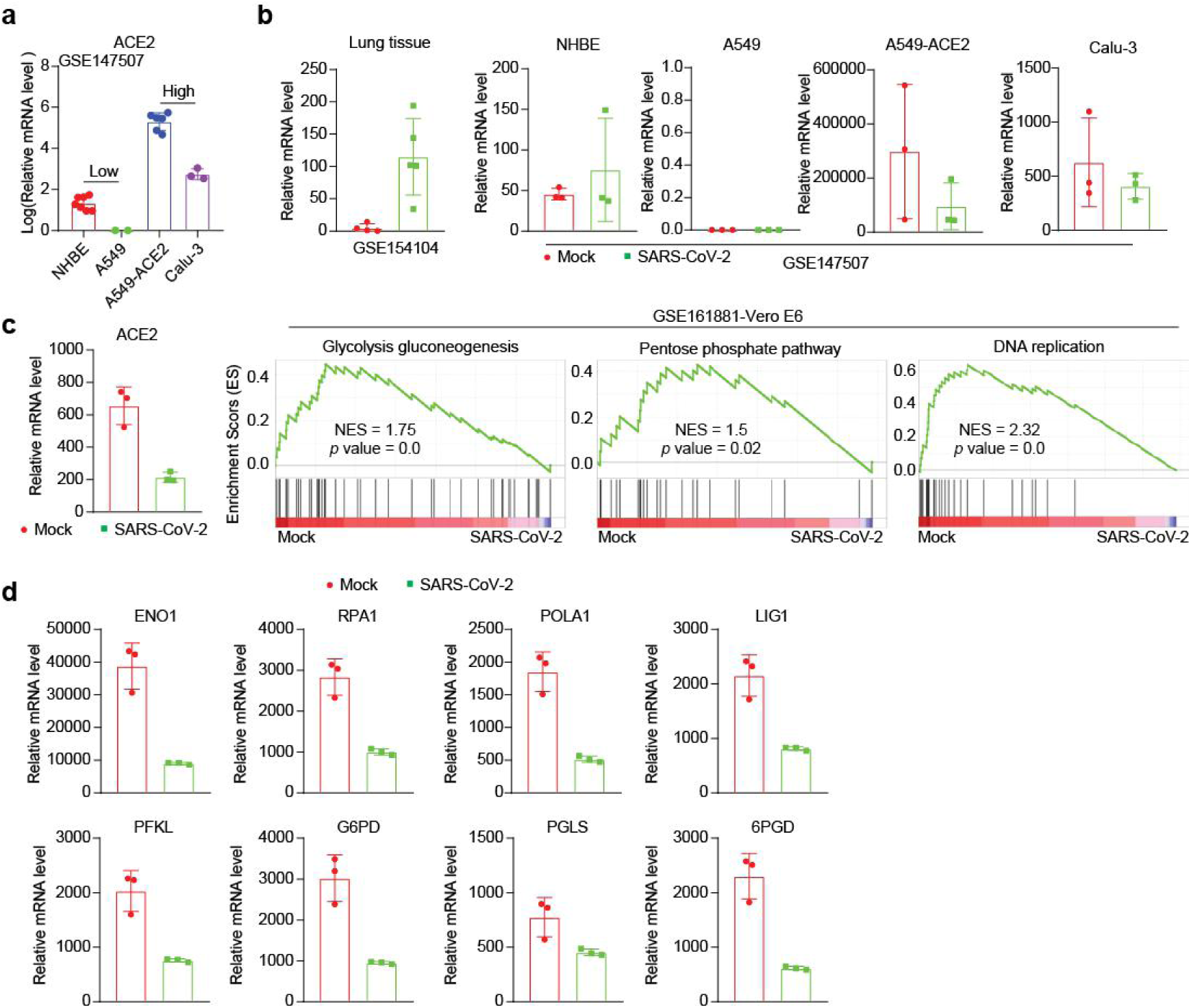
ACE2 determines how virus alters host’s metabolic balance. **a** The expression level of ACE2 in different lung cells. **b** Expression changes of ACE2 in mice lung tissue, lung normal cells, and lung cancer cells infected with SARS-CoV-2 from the GEO datasets (GSE154104 and GSE147507). **c, d** Expression changes of ACE2, GSEA pathway enrichment analyses and expression of related genes in Vero E6 cells infected with SARS-CoV-2 from the GEO datasets (GSE161881).

To explore the exactly role of ACE2 in determining host glucose metabolism, oxiPPP and DNA replication, we examined the changes of ACE2 during SARS-CoV-2 infection. Interestingly, we found that the expression changes of ACE2 after SARS-CoV-2 infection were reversed. The expression of ACE2 was suppressed after SARS-CoV-2 infection in cells with high ACE2 expression, while the expression of ACE2 was up-regulated in cells with low ACE2 expression (Fig. 3b). In order to verify the accuracy of our results again, we selected Vero E6 cells as host cells,^40^ as the expression levels of ACE2 in Vero E6 cells is almost same as that in Calu-3 cells, so Vero E6 cells are considered to be high ACE2 expressing cells.^41^ Indeed, our analysis results showed that SARS-CoV-2 infection inhibited the expression of ACE2, and down-regulated glucose metabolism, oxiPPP and DNA replication, as well as decreased the related enzymes expression (Fig. 3c and d). Therefore, our analysis showed that the cells with high ACE2 expression, which infected with SARS-CoV-2 would decreased DNA replication due to the decreased ACE2 levels. In the reported paper, we also found that there are no consistent conclusion on the expression changes of ACE2 in host with SARS-CoV-2 infection,^30,42–44^ which may be due to the differences in the expression levels of ACE2 among host cells before infection. This discovery made us realize that ACE2 not only mediates the SARS-CoV-2 infection, and also involves in the process of regulating viral nucleic acid replication. Thus, modulating the levels of ACE2 is a key factor for determining viral nucleic acid replication. Coincidentally, ACE2 is transcriptionally regulated by BRD4, and the inhibition of BRD4 by drugs (ABBV-744) inhibits the replication of SARS-CoV-2.^45^ This is an exciting result, and the discovery will provide not only a powerful boost in the fight against COVID-19, and also theoretical help for the development of more accurate methods for identifying positive infections in asymptomatic infections.

### ACE2 promotes nucleotide replication required for viral infection

As replication of many viruses is supported by enhanced aerobic glycolysis,^24,29^ we believed that SARS-CoV-2 replication in host cells (especially lung cells) is reliant upon altered glucose metabolism. This metabolism is similar to the Warburg effect well studied in cancer. Therefore, after finding that the receptor protein ACE2 mediates the infection and metabolic reprogramming of SARS-CoV-2, we further analyzed the role of ACE2 in lung cancer. The results showed that the expression of ACE2 was up-regulated in lung cancer patients compared with normal people from GEO database, suggesting that ACE2 also has a promotion role in lung cancer (Fig. 4a). After dividing healthy people and lung cancer patients from TCGA database into high and low expression groups of ACE2, we found that high expression of ACE2 promoted glucose metabolism and oxiPPP (Fig. 4b), but DNA replication were not enriched. Likewise, previously screened target genes were not affected by ACE2 expression (Fig. 4c), except for the three metabolic enzymes of oxiPPP. However, other metabolic enzymes of glucose metabolism (ADH6, ADH7, ACSS2, ADH4, ALDOA and ALDOB) were positively correlated with ACE2 expression and their expression was up-regulated in high ACE2 group (Fig. 4d). In addition, we analyzed RNA-sequencing results in Calu-3 knockdown ACE2 cells ^45^ and found that glucose metabolism and oxiPPP were also enriched, but not statistically significant (Fig. 4e), while DNA replication was not enriched at all, which may be due to the sample only have two replicates. When we analyzed the expression levels of specific genes, genes (PGK1, ENO1, and PGAM1) regulated by SARS-CoV-2 infection were not affected by ACE2 knockdown (Fig. 4f). While the expression levels of related enzymes (ADH6, ADH7, ALDOB, G6PD, and 6PGD) identified in lung cancer were down-regulated after ACE2 knockdown (Fig. 4g). These results suggest that although ACE2 promotes glucose metabolism in both viral infections and tumors, the metabolic enzymes regulated are not the unanimous. Tumor cells need more energy and materials for cell proliferation; while virus has a greater demand for nucleotides replication. The changes of metabolic enzymes in oxiPPP only indicate that tumor cells also have needs for nucleic acid replication. In short, ACE2 is important for lung cancer through reprogramming glucose metabolism not DNA replication, while, ACE2 modulates the ability of host cell’s DNA replication under SARS-CoV-2 infection condition.

**Fig. 4.**
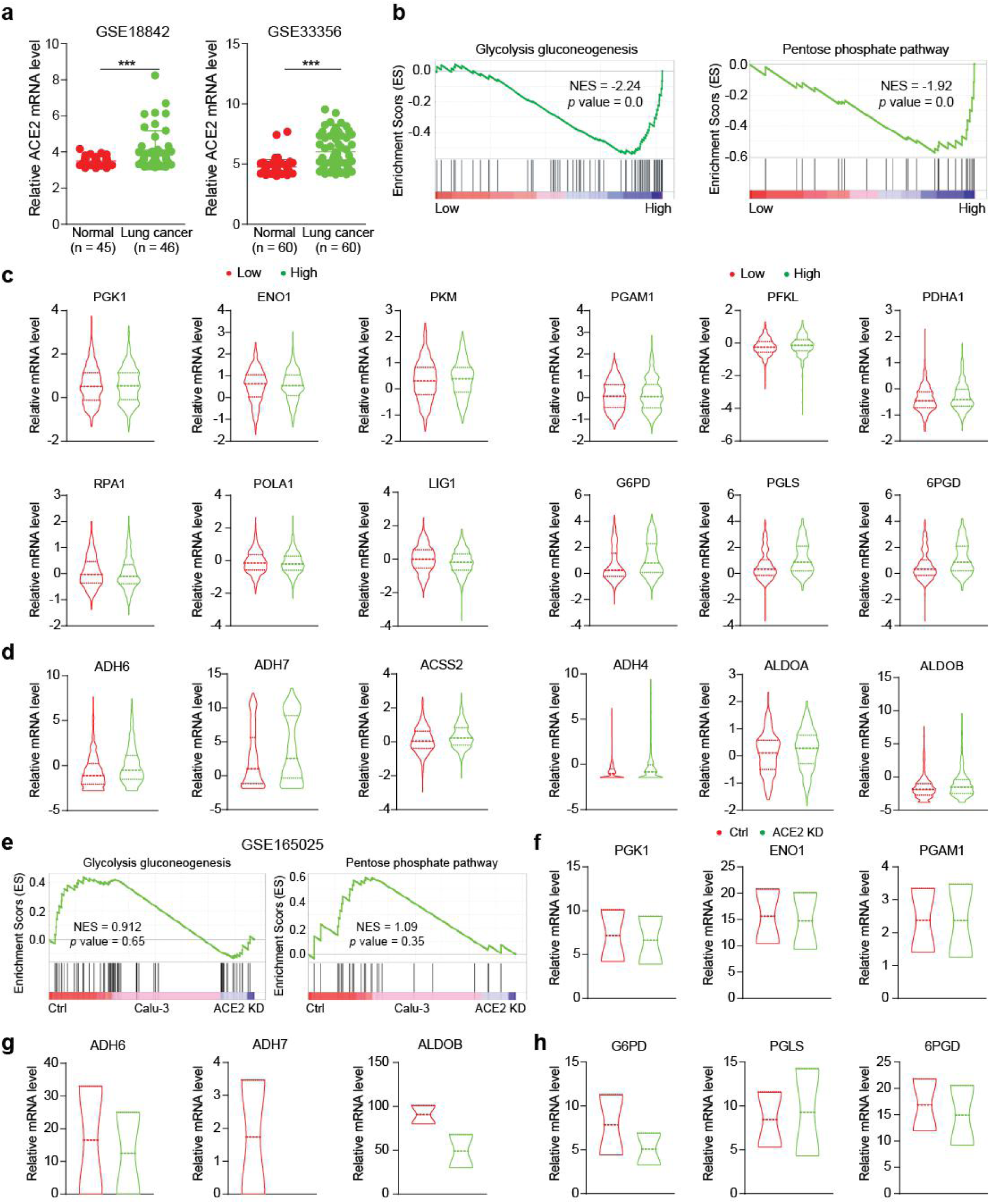
ACE2 promotes glucose metabolism in lung cancer. **a** The ACE2 mRNA expression levels between tumor and normal tissues of patients with lung cancer from GEO database (GSE18842 and GSE33356). **b** GSEA pathway enrichment analyses of ACE2 signature in healthy people and patients with lung cancer from the TCGA-LUNG datasets. **c** The expression of screened target genes in high and low ACE2 group. **d** Expression changes of glucose metabolic enzymes in high and low ACE2 group. **e** GSEA pathway enrichment analyses of ACE2 knockdown in Calu-3 cells from the GEO datasets (GSE165025). **f-h** The target genes expression in ACE2 knockdown Calu-3 cells from the GEO datasets (GSE165025).

### High expression of ACE2 increases the probability of asymptomatic infection

Different organisms have different susceptibility to SARS-CoV-2,^46^ and we found that mammalian primates have higher susceptibility compared to other animals (Fig. 5a-c). The evolutionary conservation of ACE2 makes us think that the expression level of ACE2 is of great significance to the susceptibility to SARS-CoV-2 (Fig. 5d). However, we cannot rule out the regulatory role of key sites in ACE2 for SARS-CoV-2 infection, and here we only focus on the role of ACE2 expression levels in SARS-CoV-2 infection and replication.

**Fig. 5.**
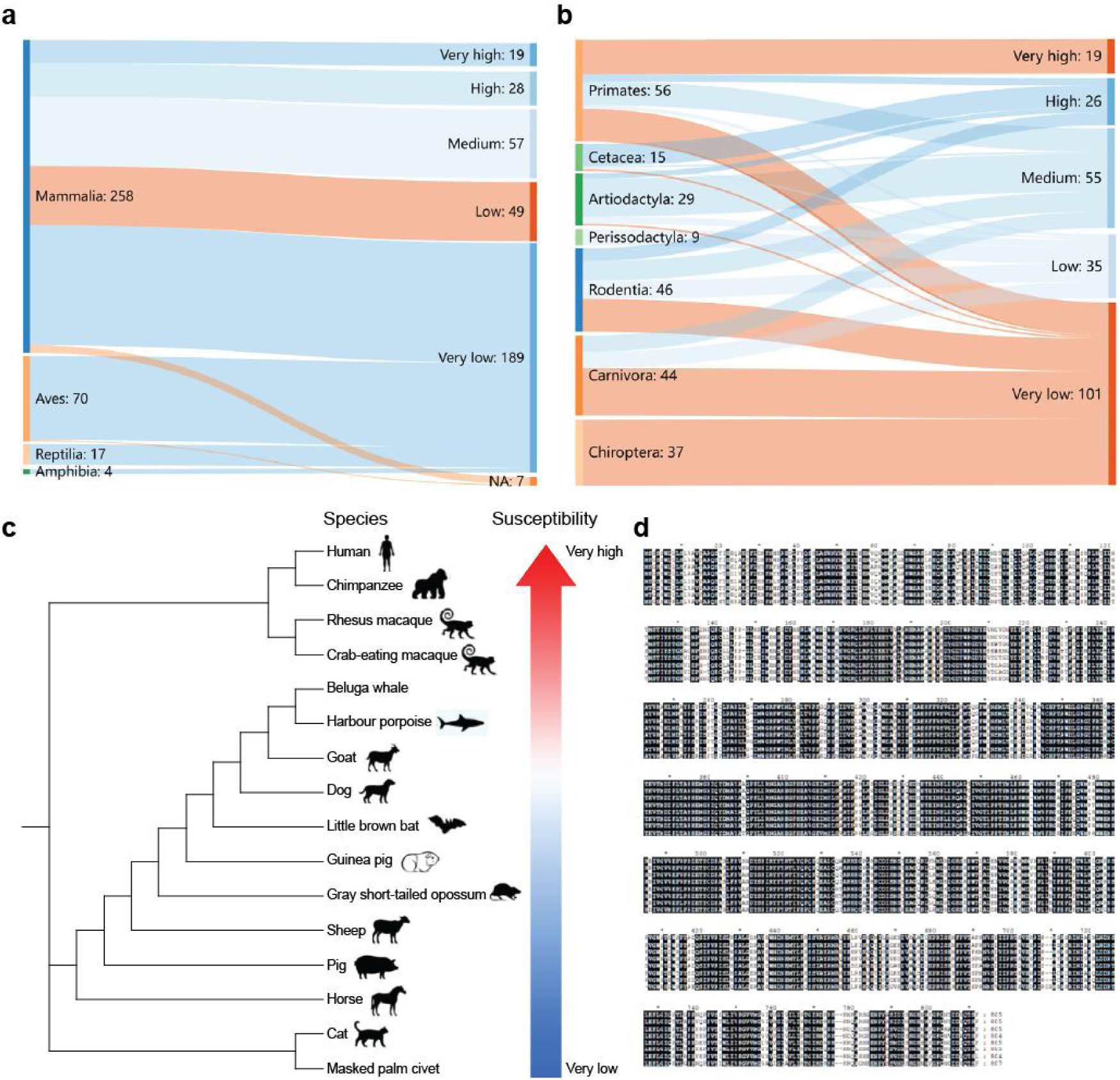
The susceptibility to SARS-CoV-2 in different species. **a, b** Comparison of susceptibility to SARS-CoV-2 across different biological classes and orders. **c, d** Phylogenetic analysis and comparison of protein sequences of ACE2.

To verify the conclusions, we first built a gender model. As we all know, the expression of ACE2 in male is higher than that in female and high ACE2 expression will make them more susceptible to infection.^47,48^ However, in the reports we collected,^49–58^ the positive rate of females was higher than that of males, and the proportion of males among asymptomatic infections was higher (Fig. 6a and b). Besides, we also collected two typical COVID-19 epidemic cases to verify our results. The first case occurred in Jinnan District, Tianjin City, China. After the first positive infection was found on January 8, 2022, government departments quickly took steps and sequenced patient samples to identify the virus as a mutant strain (Omicron: B.1.1.529). Overbearing the burden of Omicron depends crucially on the proportion of asymptomatic infections. A systematic review based on previous SARS-CoV-2 variants suggested that 40% of infections were asymptomatic^59^. Evidence suggests that the proportion of asymptomatic infections is much higher for Omicron, perhaps as high as 80-90%.^60^ As of January 23, we have collected gender information of 367 positive cases. After analysis, we found that women were consistently higher than men in cumulative cases, indicating that the virus was more easily detected in women infected (Fig. 6c). In addition, in the first three days after first confirmed case, the number of daily new positive infections among women was also higher than that of men (Fig. 6d). In second case, a student who went to school in Tianjin (Jinnan District) returned to his hometown, causing the COVID-19 epidemic in Anyang, Henan Province, China. Here, the male student was not the first case to be detected infection, but his sister who was in close contact with him was identified as a positive infection before him. Similar to Jinnan case, female were more likely to be detected for the presence of SARS-CoV-2 (Fig. 6e and f). These results suggest that women are more likely to be detected after being infected with SARS-CoV-2, which does not seem to be consistent with the reported results that men are more susceptible to infection. However, when combined with the results of our analysis, everything seems quite reasonable. It is currently recognized that the expression of ACE2 in men is higher than that in women and high ACE2 expression will make them more susceptible to infection,^61^ so men are more susceptible to SARS-CoV-2 infection.^47,48^ However, when men with high ACE2 expression are infected, glucose metabolism, oxiPPP, and DNA replication will be inhibited, which delays or blocks the replication of virus, and eventually lead to susceptible men become asymptomatic infections, even self-healing. In this way, infected people with high expression of ACE2 will prolong the detection time and increase the threat of virus transmission. Throughout the whole process, we found that it took about two weeks from the beginning to the end of the epidemic, which is also in line with the reported data.^21,22^ Besides, we also found that the sex difference disappeared at the end of the COVID-19 epidemic (Fig. 6d and f), suggesting that regulation viral replication by ACE2 occurs early in infection. In summary, we used the typical epidemiological investigation to verify our analysis results. We believe that people with high ACE2 expression are more likely to be infected with SARS-CoV-2, but at the same time these people are less likely to detect the presence of the virus at an early stage, and therefore become asymptomatic infection brings difficulties to the prevention and control of the COVID-19. However, in order to get more accurate epidemiological investigation data, we need a larger sample size to verify the findings that the host with higher ACE2 levels are prolonged the detection time or become asymptomatic infections.

**Fig. 6.**
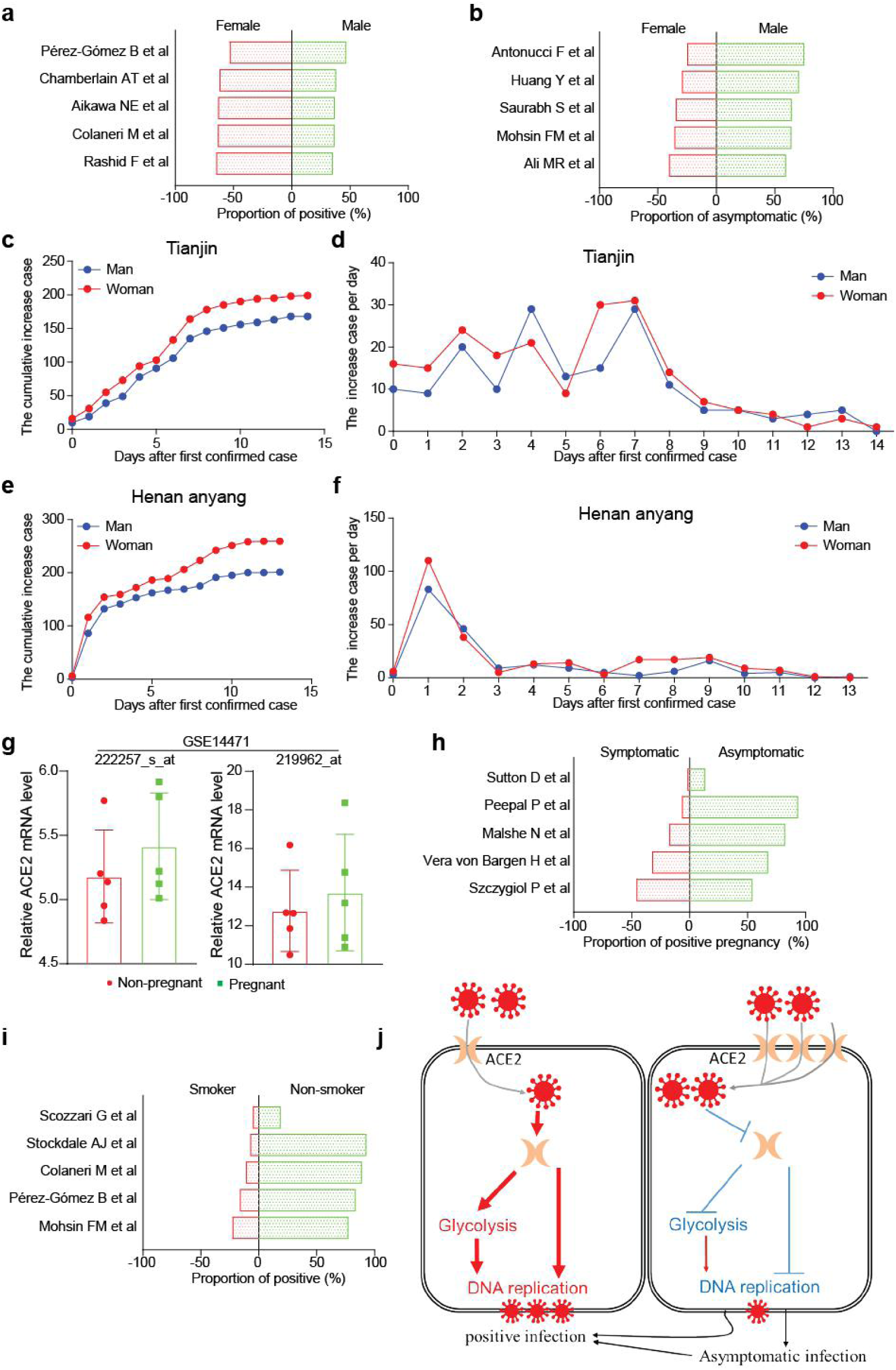
High expression of ACE2 increases the probability of asymptomatic infection. **a, b** The effect of gender on positive infection and asymptomatic infection. **c-f** Epidemiological survey and analysis in Jinnan District, Tianjin and Anyang, Henan Province, China. **g** Expression changes of ACE2 in pregnant and non-pregnant group from the GEO datasets (GSE14471). **h, i** The effects of smoking and pregnancy on positive and asymptomatic infections. **j** Schematic diagram of interaction of ACE2 to SARS-CoV-2 infection.

In addition, we also pay attention to the special group of pregnant women. We found that pregnancy lead to up-regulation of ACE2 expression (Fig. 6g). And the comparison of asymptomatic infection in pregnancy is quite high^62–66^ (Fig. 6h), which proves the important role of ACE2 in asymptomatic infection. Not only that, smoking induces ACE2 expression^67–69^ and reduces the sensitivity of infected people to detect the virus^51,53,58,70,71^ (Fig. 6i). In summary, we believe that people with high ACE2 expression are more likely to be infected with SARS-CoV-2, but these people are less likely to detect the presence of virus at an early stage, and therefore become asymptomatic infection brings difficulties to the prevention and control of COVID-19.

## CONCLUSION

The COVID-19 epidemic has been going on for more than 2 years, and the whole world has been shrouded in the shadow of SARS-CoV-2, unable to return to normal life and work. As people with certain scientific research literacy and ability, we should do our best to make certain efforts to prevent and control the epidemic. Here, by analyzing the existing data and combining with epidemiological investigations, we believe that ACE2 not only mediates the process of SARS-CoV-2 infection and invasion of the host, but also participates in the regulation of glucose metabolism to support SARS-CoV-2 replication. So, our analysis results suggested that the host with high ACE2 expression has a certain inhibitory effect on virus replication (Fig. 6j). As the saying goes: “whoever started the trouble should end it”. Collectively, we sincerely hope that these results will be known to more scientific researchers, provide help for the prevention and control of the epidemic, and develop more accurate detection methods to prevent the spread of the SARS-CoV-2. However, how SARS-CoV-2 infection altered the expression of ACE2 in the host cells with different ACE2 levels to reprogram glucose metabolism for supporting DNA replication remains unclear.

## ACKNOWLEDGMENTS

The study was supported by grants from National Nature Science Foundation of China (81973356 to C.S., 81902826 and 81672781 to S.Z.), the Fundamental Research Funds for the Central Universities of Nankai University (3206054, 91923101, 63213082, and 92122017 to C.S.), the State Key Laboratory of Drug Research (SIMM2105KF-08 to C.S.), and the National Key R&D Program of China (No. 2021YFC1712900 to S.Z.).

## AUTHOR CONTRIBUTIONS

Conceptualization, J.W. and C.S.; Methodology, J.W., and H.C.; Investigation, J.W., Y.Q., and H.S.; Writing - Original Draft, J.W., X.L., and C.S.; Writing - Review & Editing, J.W., S.Z., and C.S.; Funding Acquisition, S.Z. and C.S.; Resources, S.Z. and C.S.; Supervision, S.Z. and C.S.

## DECLARATION OF INTERESTS

The authors declare no competing interests.

## Notes

### Competing Interest Statement

The authors have declared no competing interest.

